# Polyamines and eIF5A hypusination facilitate SREBP2 translation and cholesterol synthesis to enhance enterovirus attachment and infection

**DOI:** 10.1101/2021.11.01.465941

**Authors:** Mason R. Firpo, Marine J. Petite, Natalie J. LoMascolo, Priya S. Shah, Bryan C. Mounce

**Author notes:** Corresponding author: Department of Microbiology and Immunology, Loyola University Chicago, Stritch School of Medicine, 2160 S. First Ave., Maywood, IL 60153.

## Abstract

Metabolism is key to cellular processes that underlie the ability of a virus to productively infect. Polyamines are small metabolites vital for many host cell processes including proliferation, transcription, and translation. Polyamine depletion also inhibits virus infection via diverse mechanisms, including inhibiting polymerase activity and viral translation. We showed that Coxsackievirus B3 (CVB3) attachment requires polyamines; however, the mechanism was unknown. Here, we report polyamines’ involvement in translation, through a process called hypusination, promotes expression of cholesterol synthesis genes by supporting SREBP2 translation, the master transcriptional regulator of cholesterol synthesis genes. Measuring bulk transcription, we found polyamines support expression of cholesterol synthesis genes, regulated by SREBP2. Polyamine depletion inhibits CVB3 by depleting cellular cholesterol. Exogenous cholesterol rescues CVB3 attachment, and mutant CVB3 resistant to polyamine depletion exhibits resistance to cholesterol perturbation. This study provides a novel link between polyamine and cholesterol homeostasis, a mechanism through which polyamines impact CVB3 infection.

## Introduction

Polyamines are small carbon chains with amine groups that have a positive charge at cellular pH. They play a large role within the cell and are involved in multiple cellular processes including nucleotide synthesis, DNA/RNA stability, membrane fluidity, and translation^1^. Ornithine, which is a derivative of arginine, is converted to the first polyamine putrescene by the rate limiting enzyme ornithine decarboxylase 1 (ODC1). Putrescine can be converted to spermidine and spermine via their respective synthases. Difluoromethylornithine (DFMO) is an FDA approved drug for trypanosomiasis and is an irreversible, competitive inhibitor of ODC1^2^. One key way polyamines impact cells is through translation, specifically through a process called hypusination^3^. Spermidine is covalently attached to eukaryotic initiation factor 5A (eIF5A) at lysine 50 by the protein deoxyhypusine synthase (DHPS) to form deoxyhypusine-eIF5A. Deoxyhypusine hydroxylase (DOHH) then adds a hydroxide in the second and final step to make hypusine-eIF5A. Hypusine-eIF5A plays a vital role in mRNA translation, ribosome function, and cell proliferation. The precise mechanisms by which hypusine-eIF5A promotes translation remain to be fully understood. However, certain amino acid motifs cause ribosomal pausing, and hypusine-eIF5A is required for the ribosome to translate through these motifs^4^. Poly-proline tracts have also been shown to require hypusine-eIF5A^5^. When the unhypusinated form of eIF5A is present, it cannot alleviate ribosomal pausing. The requirement for hypusine-eIF5A for cellular proliferation has made it an attractive target for the development of anti-cancer drugs. One such drug is the spermidine analog N1-guanyl-1,7-diaminoheptane (GC7). GC7 inhibits DHPS by directly binding to the active site and prevents spermidine from being attached to eIF5A^6^. Additionally, deferiprone (DEF) is an iron-chelator which has broad effects on the cell and also inhibits DOHH^7^.

Polyamines have been found to be important for multiple RNA viruses through different mechanisms, and diverse RNA viruses are sensitive to DFMO-mediated polyamine depletion^8–11^. Interestingly, the enterovirus Coxsackievirus B3 (CVB3) develops mutations within its proteases as well as the capsid protein VP3 when polyamine synthesis is inhibited, suggesting a role for polyamines in protease activity and cellular attachment for enteroviruses^12–14^. Enteroviruses are small, non-enveloped, positive sense single-stranded RNA viruses a part of the picornavirus family that can cause a range of diseases from the mild cold to flaccid paralysis and dilated cardiomyopathy (DCM)^15,16^. Coxsackievirus B3 (CVB3) is a member of the enterovirus genus and is well known for its ability to infect and persist in the heart and cause DCM^17,18^. About 50% of DCM patients have CVB3 reactive antibodies and the only cure for DCM is a heart transplant^16,19^. There are currently no FDA drugs approved to treat CVB3 infection. However, we previously found that inhibiting polyamines is broadly antiviral and inhibits CVB3 infection and binding^12,13,20^.

Another key metabolite required for CVB3 infection is cholesterol. Cholesterol is important for maintaining cellular membrane integrity and fluidity as well as lipid raft formation. Removal of cholesterol from the plasma membrane blocks poliovirus entry and EV-11 entry by preventing lipid raft formation^21,22^. Inhibiting cholesterol homeostasis significantly inhibits CVB3 replication^23^. The first step in cholesterol synthesis is the conversion of Acetyl-CoA to HMG-CoA by HMGC-CoA synthase (HMGCS). HMG-CoA is then converted to mevalonic acid by the rate limiting enzyme HMG-CoA reductase (HMGCR). After 27 more reactions, the end product of cholesterol is made^24^. The majority of these genes, including low-density lipoprotein receptor (LDLR) is under the transcriptional control of sterol regulatory element binding protein 2 (SREBP2), which binds to sterol regulatory elements (SREs) in the promoter of target genes^25^. Upon depletion of cholesterol, the ER resident, multipass transmembrane protein, SREBP2 is translocated to the Golgi. Within the Golgi it gets cleaved by the proteases S1P and S2P generating the active N-terminal portion of the protein. The active SREBP2 then re-locates to the nucleus where it promotes the transcription of sterol synthesis genes^26^. To date, no link has been established between cholesterol synthesis and polyamines; however, mice overexpressing the polyamine catabolic enzyme spermidine-spermine acetyltransferase (SAT1) and rats treated with DFMO exhibited lower serum cholesterol levels^27,28^, suggesting that polyamines may facilitate cholesterol synthesis.

Using a transcriptomic approach, we identified several pathways modulated by polyamines that likely impact enterovirus infection. We found that cholesterol synthesis genes were enriched in this analysis and hypothesized that polyamines may impact cholesterol synthesis and virus attachment. Here we describe a novel link between polyamines and cholesterol synthesis through the polyamine-dependent translation of SREBP2. We find that inhibition of polyamine synthesis or specific inhibition of hypusination leads to reductions in SREBP2 translation, activity, and downstream gene expression. This culminates in reduced cellular cholesterol and, in turn, reduced viral attachment and replication. These data connect previously unrelated metabolic pathways in the cell and identify cellular cholesterol depletion as an important effect of polyamines on virus replication, with important implications for both virus infection and cellular metabolic status.

## Results

### Inhibition of polyamine synthesis inhibits CVB3 binding and is rescued with exogenous polyamines

We previously found that inhibition of polyamine synthesis by the suicide inhibitor DFMO, significantly decreases CVB3 binding to cells compared to untreated cells^13^. To confirm this phenotype, we treated Vero cells for 4 days with increasing doses of DFMO to deplete cellular polyamines. We then added CVB3 directly to cells on ice for 5 minutes. Virus was then washed off followed by agar overly, in media containing polyamines. Thus, in these assays, polyamines are depleted only for attachment. Plaques generated from successful attachment and entry were allowed to form and developed two days later, and bound virus was enumerated by counting these plaques (Fig. 1A). We found that DFMO significantly reduced bound virus in a dose-dependent manner (Fig 1B), in agreement with prior work and corresponding to a decrease in cellular polyamines, as measured by thin layer chromatography (Fig. 1C). To determine if this polyamine-dependent attachment phenotype relied on cellular factors, we treated cells with DFMO to deplete polyamines and subsequently replenished the polyamines (putrescine, spermidine, and spermine) in an equimolar concentration. Adding polyamines to the cells at the time of infection did not rescue viral attachment, nor did addition 4h prior to attachment. However, when polyamines were added 16h prior to infection, we observed a full rescue in CVB3 attachment. These data suggest that polyamines rescue viral attachment but rely on an extended incubation period, perhaps because cellular synthesis of attachment factors required an extended time.

**Figure 1.**
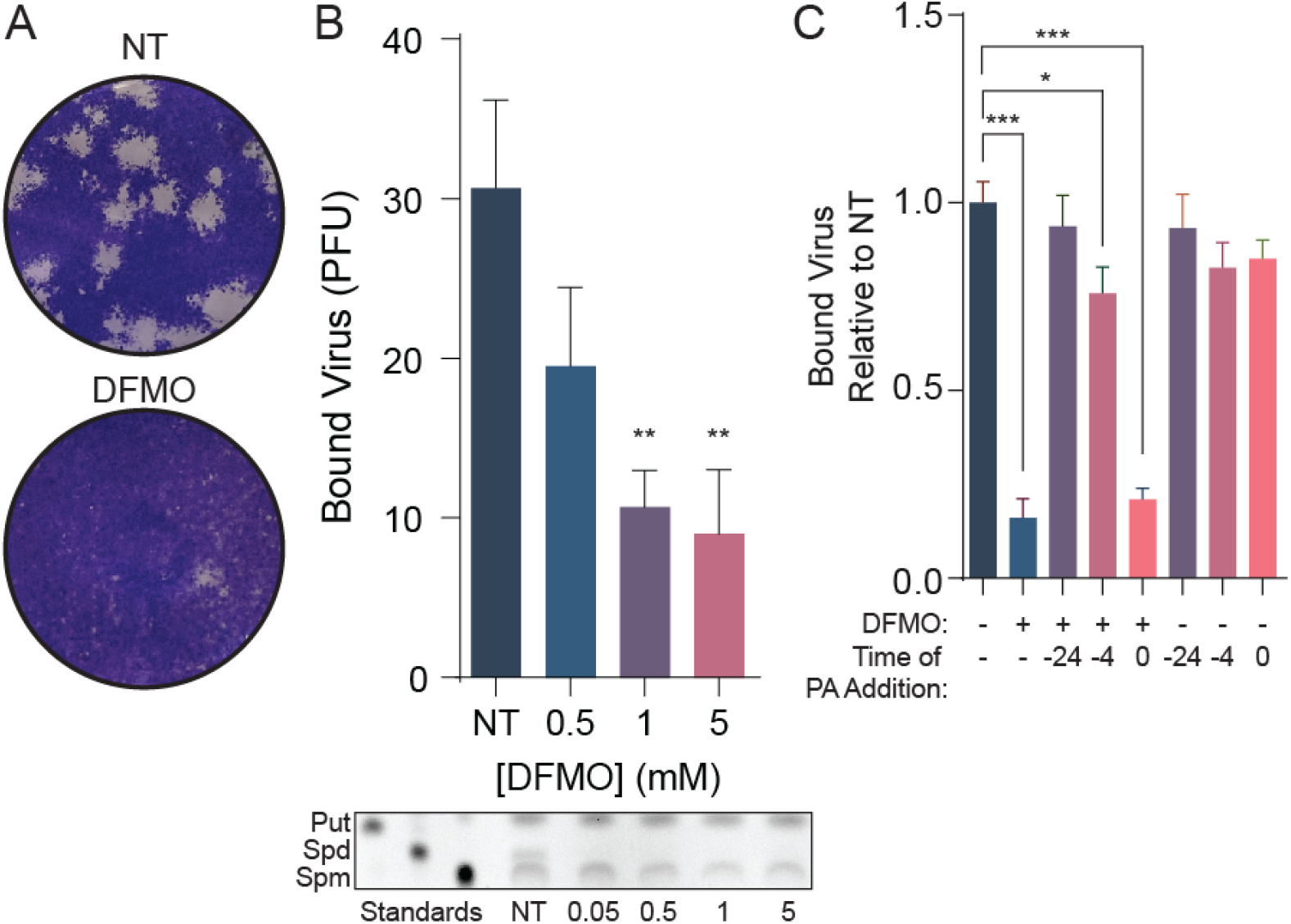
CVB3 requires polyamines for attachment. (A) Representative plaques from B. (B) (Top) Quantification of plaques formed from a CVB3 binding assay with DFMO treated Vero cells. (Bottom) Thin layer chromatography of Huh7 cells treated with increasing doses of DFMO. (C) Vero cells were treated with DFMO for 96 hours then treated with 10 μM equimolar ratio of polyamines at the indicated times before infection. \**p* < 0.05, \*\**p* < 0.01, and \*\*\**p* < 0.001 by the Student’s *t* test. Data from at least three independent experiments.

### Polyamine depletion decreases expression of genes in the cholesterol synthesis pathway

To better understand the effects polyamine depletion has on cells and if we could identify polyamine-modulated cellular factors involved in viral attachment, we performed RNA-sequencing on untreated and polyamine-depleted cells. Huh7 cells were left untreated or depleted of polyamines with 1 mM DFMO for 96 h, at which time RNA was extracted and analyzed by Illumina paired-end reading. After the alignment of reads against human genome, a differential gene expression analysis was conducted to identify significant changes in expression. To uncover the underlying biological processes that may link these pathways, gene set enrichment analysis (GSEA) was performed. This showed multiple metabolic pathways that were significantly enriched for decreased gene expression by polyamine depletion including alcohol metabolism, cholesterol metabolism, and cellular response to a chemical (Fig 2A). Cholesterol plays a large role in membrane stability and fluidity which could account for the differences seen in the GSEA analysis. In order to investigate specific cholesterol genes affected by DFMO, cholesterol genes involved directly in cholesterol synthesis were overlayed in a Volcano plot (Fig 2B). Multiple genes were down regulated including HMGCS1, HMGCR, MVK, and MVD. Importantly, several polyamine metabolic genes, including SAT1 and OAZ1 exhibited reduced expression, consistent with their role in inhibiting polyamine synthesis. We next explored how these changes caused by DFMO related to changes caused by CVB3 infection. Again, differential gene expression analysis was performed on CVB3-infected and mock-treated cells. Leading edge genes from the “Cholesterol Homeostasis” category were visualized by heat map for CVB3-infected and DFMO-treated samples (Fig 2C). Interestingly, CVB3 infection increased expression of many cholesterol biosynthesis genes that are downregulated by DFMO, suggesting CVB3 has mechanisms to promote this pro-viral pathway. Thus, the transcriptomic analysis of polyamine depleted and virus-infected cells revealed that cellular metabolic processes and, specifically, cholesterol biosynthesis (Fig 2D) may function in a pro-viral manner.

**Figure 2.**
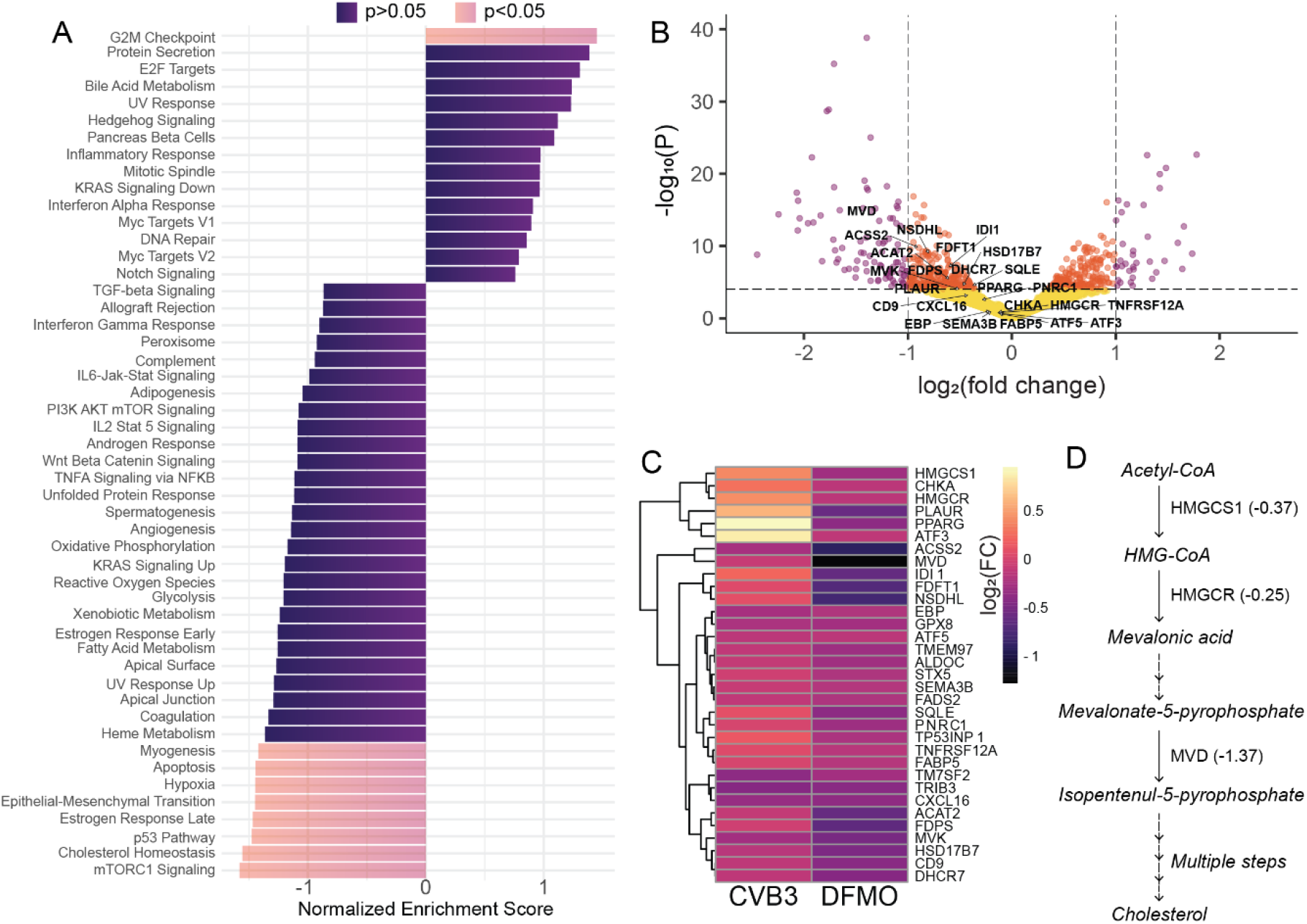
Transcriptomic analysis of DFMO treated cells. Huh7 cells were treated with 1mM DFMO for 96h and subjected to RNA sequencing. Results are based on duplicates. (A) GSEA was conducted on genes differentially expressed by 1mM DFMO treated Huh7 versus untreated Huh7. The top positively and negatively enriched Hallmark pathways were plotted (p adj < 0.05, purple, p.adj >0.05, blue). (B) Volcano plot indicating log2FoldChange for genes from differential gene expression analysis comparing DFMO treated cells relative to untreated cells. Significant changes in gene expression are plotted in purple for genes : p.adj < 0.05, log2FC > 1, in orange for p_adj_<0.05, log_2_FC<1 and in yellow for p_adj_ >0.05, log_2_FC<1. P-values were adjusted for false discovery rate using Benjamini Hochberg method. (C) Genes from the leading edge of cholesterol homeostasis pathway were subjected to hierarchical clustering for both conditions DFMO treated or CVB3 infected cells relative to untreated cells. Log_2_ fold change from target genes are displayed as a heat map. (D) Cholesterol synthesis pathway with the representation of down-regulated genes HMGCS1, HMGCR and MVD.

### Polyamine depletion decreases cholesterol gene expression, protein levels, and intracellular cholesterol

To test polyamines’ effect on the transcription of cholesterol synthesis genes and confirm the RNA-seq data, cells were treated for four days with increasing concentrations of DFMO. RNA was then extracted and RT-qPCR was performed using optimized and specific primers. HMGCR, HMGCS, and MVD showed moderate but significant reductions in expression (Fig. 3A-C), aligning with the RNA-sequencing data. To determine if this reduction in transcription affected protein synthesis, we examined total cellular levels of HMGCR and MVD by western blot. Both proteins showed reduced levels with polyamine depletion (Fig. 3D). Finally, to determine if reduction of transcription and translation of cholesterol synthesis proteins affected intracellular cholesterol, cells were treated with DFMO for 96h, and the total amount of cellular cholesterol was measured via a luciferase-based cholesterol assay. We found that total cellular cholesterol was significantly reduced with DFMO treatment, consistent with a decrease in expression of cholesterol synthesis genes (Fig. 3E). Thus, cellular cholesterol synthesis relies on polyamines through the expression of cholesterol synthetic proteins.

**Figure 3.**
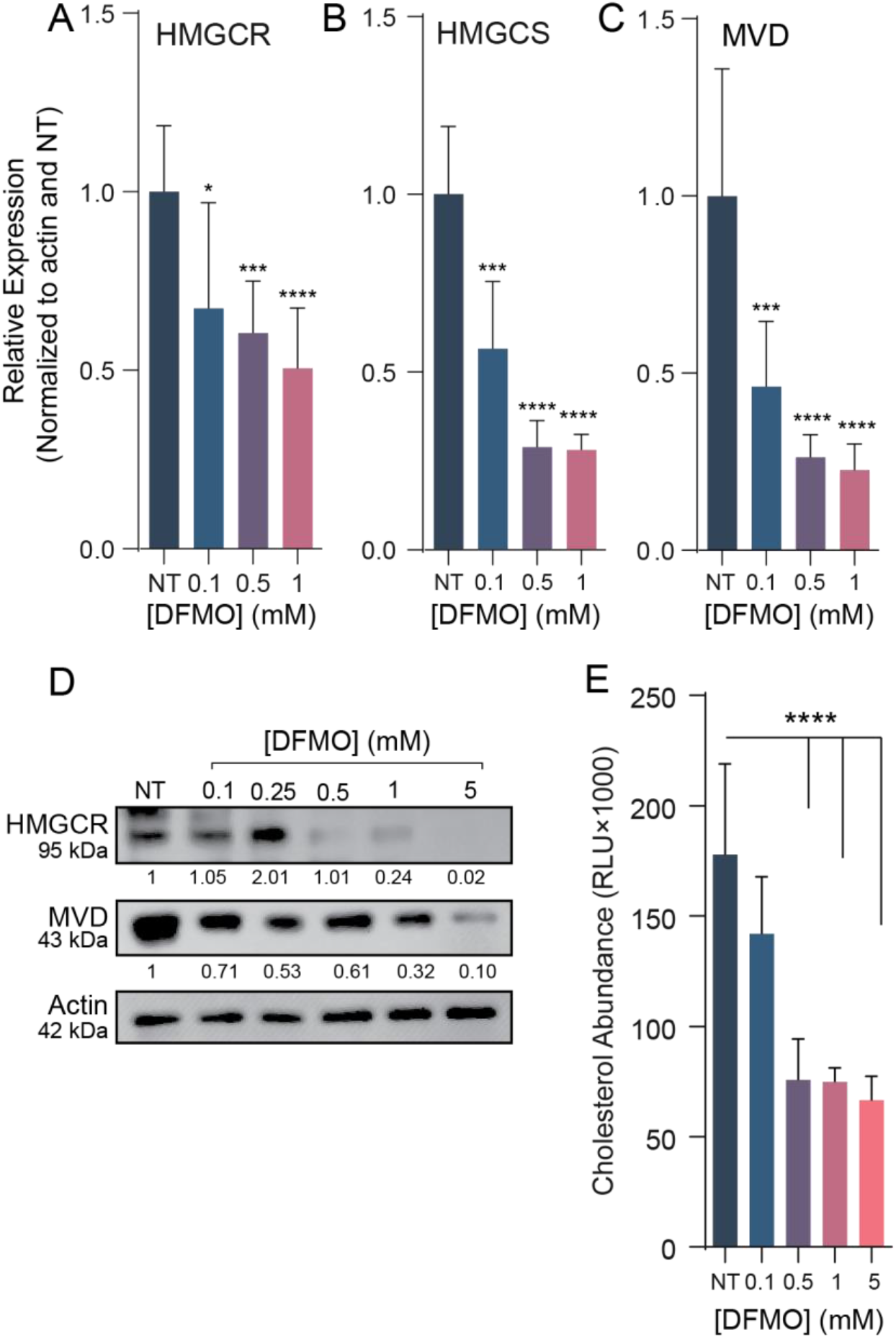
Depletion of polyamines decreases cholesterol synthesis gene expression, translation, and intracellular abundance. (A-C) Normalized qPCR of cholesterol synthesis genes relative to actin expression in Huh7 cells treated with DFMO. (D) Western blot of Huh7 cells treated with increasing doses of DFMO. Actin was used as a loading control. (E) Intracellular cholesterol abundance in DFMO treated Huh7 cells. \**p* < 0.05, \*\**p* < 0.01, \*\*\**p* < 0.001, and \*\*\*\**p* < 0.0001 by the Student’s *t* test. Data from at least three independent experiments.

### Polyamine-dependent SREBP2 translation but not transcription facilitates transcriptional activity at cholesterol gene promoters

Having observed that an array of cholesterol synthetic enzymes were reduced in transcription and translation by polyamine depletion, we considered that polyamines may affect a master regulator of their expression, rather than on each gene individually. We hypothesized that polyamines were affecting the transcription factor sterol regulatory binding protein 2 (SREBP2), one such regulator. SREBP2 is a multipass transmembrane protein found within the ER. When cholesterol levels are low within the cell, SREBP2 relocates to the Golgi where it is cleaved by S1P and S2P. The N-terminus of SREBP2 then relocates to the nucleus where it binds to sterol regulatory elements (SRE) to increase transcription of cholesterol synthesis genes. To test the impact polyamines have on SREBP2, Huh7 cells were treated with increasing doses of DFMO followed by qPCR (Fig. 4A). Unlike the cholesterol synthetic genes, we observed no significant change in SREBP2 transcripts. However, examining SREBP2 protein levels by western blot revealed that polyamine depletion caused a reduction of SREBP2 protein (Fig. 4B). To determine if this reduction of SREBP2 translation was sufficient to impact the expression of cholesterol synthesis genes, we measured the activity of SREBP2 binding to its promoter, the SRE. We transfected cells with or without polyamines with a construct encoding firefly luciferase driven by distinct cellular SREs. We used the HMGCS SRE, the LDLR SRE, and a generalized SRE created with the SRE consensus sequence. Additionally, we transfected renilla luciferase to control for effects of polyamine depletion on luciferase translation. When we measured SRE activity, we noted a significant reduction in activity in polyamine depleted cells for all SREs tested, suggesting that polyamine depletion impacts SRE promoter activity, likely due to a reduction in SREBP2 translation. Thus, polyamines facilitate translation and activity but not transcription of SREBP2.

**Figure 4.**
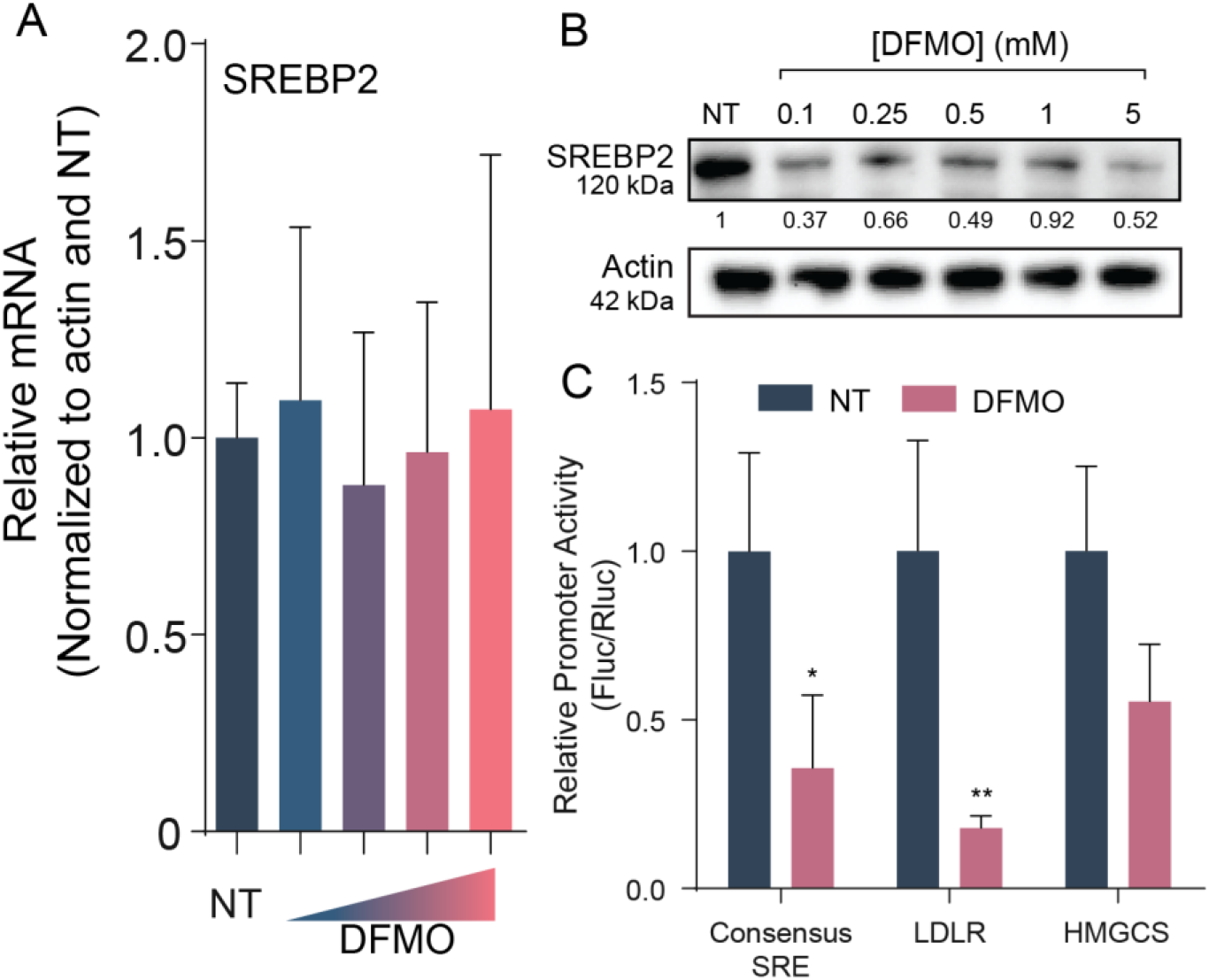
SREBP2 translation is dependent on polyamines. (A) Normalized qPCR of SREBP2 with increasing doses of DFMO (0.1, 0.5, 1, 5mM) relative to actin expression in Huh7 cells. (B) Western blot of SREBP2 in DFMO treated Huh7 cells. Actin was used as a loading control. (C) Huh7 cells were treated with 1mM DFMO for 96h followed by transfection with the promoter luciferase constructs and siCheck. Results are normalized to NT and are relative to renilla activity. \**p* < 0.05 and \*\**p* < 0.01 by two-way ANOVA. Data from at least three independent experiments.

### Polyamine-dependent eIF5A hypusination supports SREBP2 translation and cholesterol synthesis

A well-described mechanism by which polyamines support cellular translation is through the post-translational modification of eIF5A, in which spermidine is conjugated and hydroxylated, forming a unique amino acid called hypusine (Fig. 5A). While the precise mechanism(s) by which hypusinated eIF5A facilitates translation remain incompletely understood, it is known that a subset of cellular proteins rely on this enzyme for efficient translation. To determine if SREBP2 is included in this subset, we treated cells with inhibitors of the hypusination pathway. Inhibition of the deoxyhypusine synthase (DHPS) inhibitor GC7 resulted in a dose-dependent reduction in SREBP2 protein levels, concomitant with a reduction in cellular hypusinated eIF5A (Fig 5B). In concurrence with this, treatment with GC7 resulted in significant reduction of SRE promoter activity (Fig. 5C). Additionally, we observed a reduction in HMGCR protein levels with increasing GC7 or deferiprone treatment (Fig. 5D). To confirm that these changes in SREBP2 and cholesterol synthesis gene expression affected cellular cholesterol, we again measured total cellular cholesterol in cells treated with increasing doses of GC7 (Fig. 5E) or the deoxyhypusine hydroxylase (DOHH) inhibitor deferiprone (DEF) (Fig. 5F). Similar to our results with DFMO, we observed a significant, dose-dependent reduction in cellular cholesterol when hypusination was inhibited, suggesting that polyamines facilitate cholesterol synthesis through hypusination. Finally, to confirm that hypusinated eIF5A contributes to viral attachment, as we see with DFMO, we treated cells with increasing doses of GC7, performed an attachment assay, and counted attached viruses (Fig. 5G). We observed a dose-dependent decrease of viral attachment with GC7 treatment, suggesting hypusinated eIF5A facilitates viral attachment.

**Figure 5.**
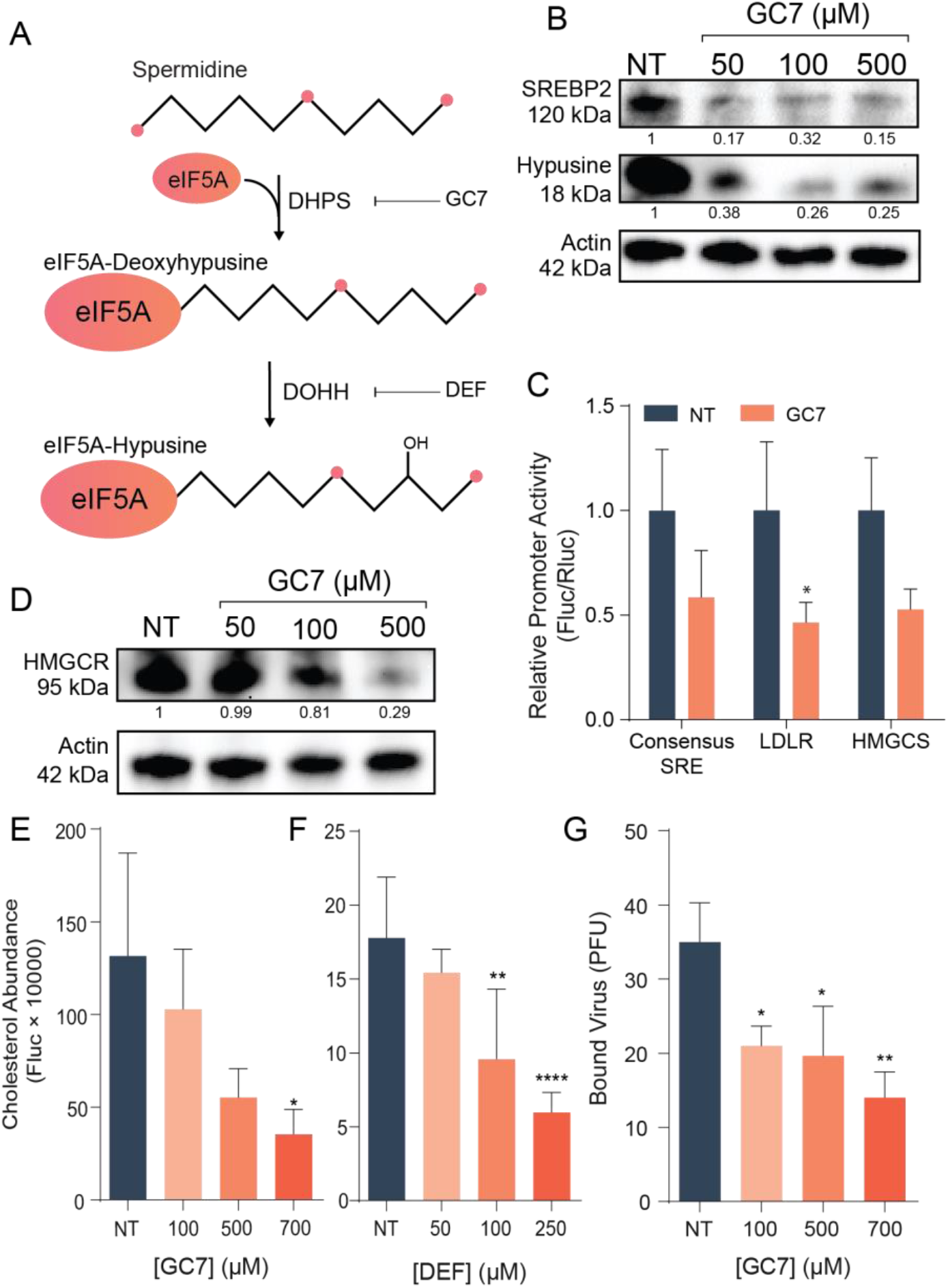
Hypusination supports cholesterol synthesis. (A) Pathway of eIF5A hypusination and its inhibition. (B) Western blot of Huh7 cells treated with increasing doses of GC7 probed for SREBP2 and hypusine-eIF5a. Actin was used as a loading control. (C) Huh7 cells treated with 500 μM GC7 for 24h followed by transfection with the promoter luciferase constructs and siCheck as a transfection control. Results were normalized to NT and are relative to renilla activity from siCheck. Two-way ANOVA was used to analyze statistical significance. (D) Western blot of Huh7 cells treated with increasing doses of GC7 and probed for HMGCR and actin was used as the loading control. (E-F) Intracellular cholesterol abundance of GC7 (E) and DEF (F) treated Huh7 cells. Y-axis is Fluc value x10000. (G) Number of CVB3 plaque forming units bound to GC7 treated Vero cells. \**p* < 0.05, \*\**p* < 0.01, \*\*\**p* < 0.001, and \*\*\*\**p* < 0.0001 by the Student’s *t* test. Data from at least three independent experiments.

### Exogenous cholesterol rescues CVB3 attachment to polyamine depleted cells and replication

Viruses require diverse cellular factors to mediate attachment and entry, including cholesterol. To determine if cholesterol is a key polyamine-modulated molecule in this process, we attempted to rescue viral attachment and replication by adding cholesterol to cells exogenously. Cells depleted of polyamines were treated with increasing doses of cholesterol overnight, followed by washing away excess cholesterol and performing a binding assay as before. When plaques were revealed, we found that cholesterol significantly rescued CVB3 binding in a dose dependent manner (Fig. 6A). To test if this rescue was due to a direct interaction of CVB3 and cholesterol or if cholesterol had to be incorporated into cellular membranes, cholesterol was added to cells at 22 h, 4 h, and 0 h before CVB3 binding. Only cholesterol added 22 h and 4 h before binding was able to significantly rescue CVB3 binding to polyamine depleted cells (Fig. 6B), suggesting that viral attachment requires cellular cholesterol incorporation. Finally, to determine if cholesterol could rescue virus replication in polyamine-depleted cells, we measured viral titers in DFMO- and cholesterol-treated cells. DFMO significantly reduced viral titers, and treatment with exogenous cholesterol significantly increased these titers, though not to untreated levels, highlighting that polyamines affect cholesterol to support viral replication but also that polyamines play multiple roles in infection.

**Figure 6.**
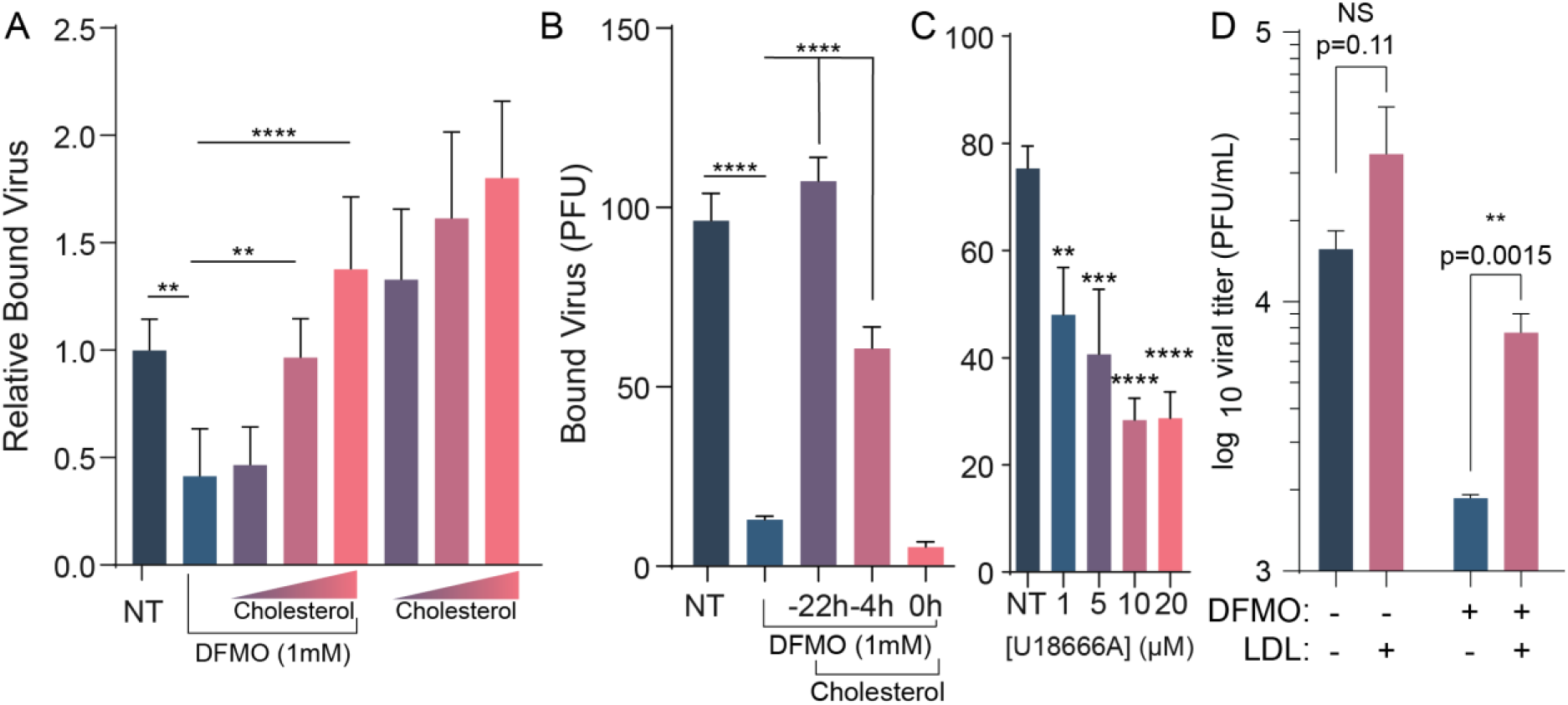
Addition of cholesterol rescues CVB3 attachment and replication in polyamine depleted cells. (A) Binding assay of Vero cells were left untreated or treated with DFMO for 96 h. After 72 h, cells were treated with 100, 250, or 500 µg/mL for 16-24h. (B) Binding assay of DFMO treated Vero cells treated with 500 µg/mL cholesterol for the indicated times. (C) Binding assay of Vero cells treated with increasing concentrations of U1866A. (D) Huh7 cells were left untreated or treated with DFMO for 96 h. After 72 h, the cells were treated with 100 µg/mL LDL for 24 h followed by infection with CVB3 at an MOI of 0.1. Virus was collected after 24 h and the PFU/mL was quantified via titration. \**p* < 0.05, \*\**p* < 0.01, \*\*\**p* < 0.001, and \*\*\*\**p* < 0.0001 by the Student’s *t* test. Data from at least three independent experiments.

**Figure 7.**
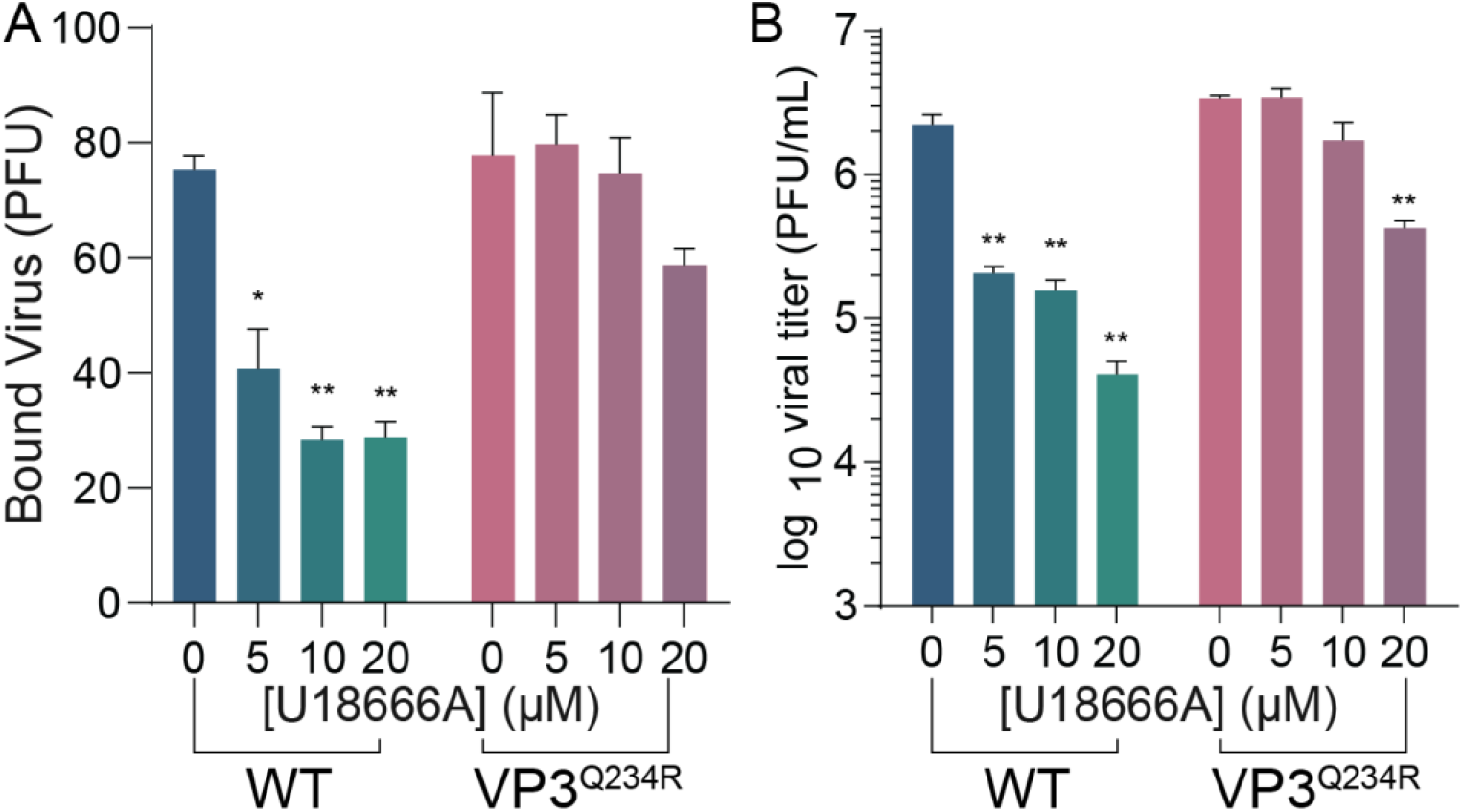
CVB3 VP3^Q234R^ mutant is resistant to reduction of cytoplasmic cholesterol. (A) Huh7 cells were treated with increasing doses of U18666A for 16h prior to binding WT and VP3^Q234R^ mutant CVB3. Bound virus was enumerated by counting plaques indicative of attached virus. (B) Cells were treated as in (A) and infected at MOI 0.1 with WT and VP3^Q234R^ mutant CVB3. Viral titers were determined at 24 hpi. *p<0.05, **p<0.01 via Student’s *t* test from three independent experiments.

**Figure 8.**
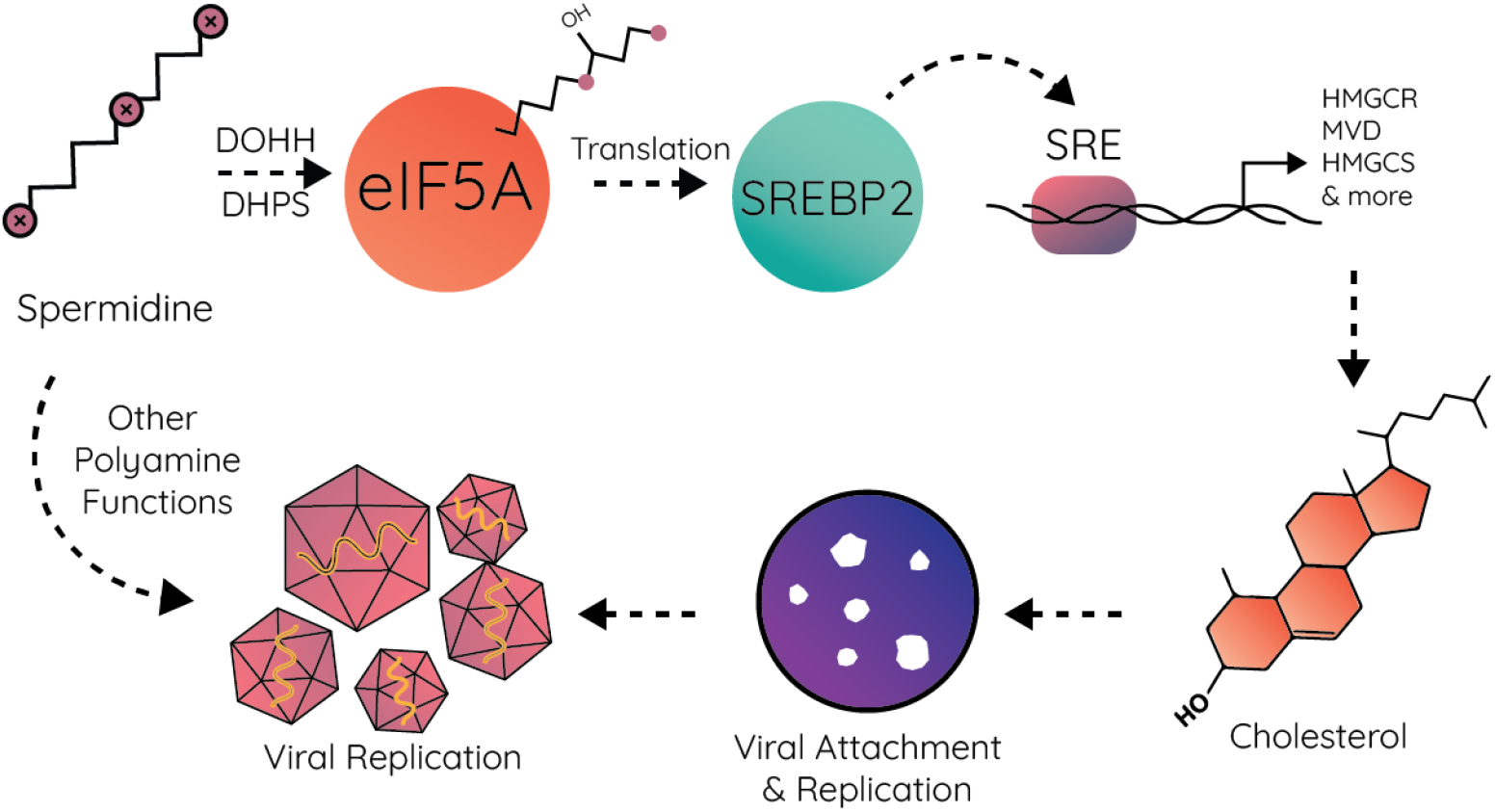
Working model. Polyamines facilitate hypusination of eIF5A, which promotes SREBP2 translation. SREBP2 binds to SREs on the promoters of cholesterol synthesis genes, leading to their expression and cholesterol synthesis. Cellular cholesterol synthesis enhances CVB3 attachment and, subsequently viral replication.

We previously described a viral mutant that exhibits enhanced viral attachment in polyamine depleted cells via mutation of VP3 at position 234. This mutant was found when we passaged virus in DFMO-treated cells, suggesting that CVB3 harboring this mutation may be resistant to polyamine depletion by overcoming a block in attachment. To determine if this block was cholesterol, we considered viral attachment in cells where cholesterol transport is impaired. The inhibitor U18666a, an NPC1 inhibitor, impacts cellular transport of cholesterol and reduces plasma membrane levels. We found that treatment of cells with U18666a reduced CVB3 replication in a dose-dependent manner, as well as attachment. We next considered that CVB3 VP3^Q234R^ may be resistant to this inhibitor. When we performed an attachment assay on U18666a treated cells using CVB3 VP3^Q234R^, we observed a modest reduction in viral attachment compared to WT CVB3, suggesting that the mutant may overcome polyamine depletion via bypassing the need for cholesterol. Additionally, we measured viral titers in U18666a-treated cells infected with CVB3 VP3^Q234R^ and observed titers significantly higher than WT CVB3. Together, these data suggest that polyamines support cholesterol synthesis to promote viral attachment and that CVB3 is able to overcome polyamine depletion by mutation of VP3, which bypasses the need for cholesterol in viral attachment.

## Discussion

Polyamines function in diverse ways within the cell, and their connections to distinct metabolic pathways are still being discovered. Here, we establish a novel connection between polyamines and cholesterol synthesis. Prior work has described how animal models treated with DFMO exhibited altered serum lipid profiles, though the molecular mechanisms remained unexplored. We found that polyamines support intracellular cholesterol levels through the hypusination of eIF5A, which is required for translation of SREBP2, a key regulator in cholesterol homeostasis. This results in reduced SREBP2 protein levels when cells are treated with DFMO or when hypusination is specifically inhibited. The reduction of SREBP2 causes a significant reduction of sterol synthesis gene expression, culminating in reduced cellular cholesterol. This polyamine-mediated cholesterol depletion also effects CVB3’s ability to attach to cells, and this binding is rescued when cholesterol is added to cells.

The process of hypusination has been well studied and defined, though the precise mechanisms by which hypusination of eIF5A supports translation remains to be fully elucidated. Hypusine-eIF5A is required for cellular proliferation and translation of “hard-to-read” sequences, such as poly-proline motifs. However, the specific genes that require hypusine-eIF5A are not well known. Here we have found that SREBP2 requires hypusine-eIF5A to be translated. While HMGCR does not have any poly-prolines, it does have a 3 amino acid motif, GGT, at position 807, that has been shown to be require hypusine-eIF5A^4^. SREBP2 on the other hand has multiple poly-proline repeats as well as other 3 amino acid motifs shown to require hypusine-eIF5A. Furthermore, a study found hypusine-eIF5A is involved in cotranslation translocation of proteins into the ER, where SREBP2 is found^29^. Our findings demonstrate another level of regulation of the cholesterol synthesis pathway, translational regulation of SREBP2. Since polyamines are also vital for cellular proliferation, a lack of polyamines will prevent crucial proliferation genes from being translated.

The roles of polyamines in viral infection are diverse and appear to be distinct for different viral families. In the case of enteroviruses, like CVB3, prior work showed that polyamines facilitate cellular attachment and protease activity. However, the mechanism(s) by which polyamines promote these activities was unclear. Cholesterol is a key molecule in enterovirus attachment, and its association with lipid rafts has been demonstrated to facilitate CVB3 engagement with its receptor (Coxsackie- and adenovirus receptor, CAR). For many viruses, cholesterol and lipids promote not only entry, but also viral replication, either through the formation of viral replication compartments on specific cellular members or through the hydrolysis of lipids to release energy for replication. While polyamines have previously been described to facilitate viral genome replication for both chikungunya virus and Ebolavirus, it remains to be determined if this phenotype could be through the synthesis of cellular cholesterol. Prior work highlighted effects of polyamines on viral proteins, such as the polymerase, but our work highlights an indirect effect on virus infection, specifically through cholesterol synthesis.

Although we show that SREBP2 relies on polyamines for translation through hypusinated eIF5A, regulation of SREBP2 and its activity is complex. Polyamines play a wide variety of roles within the cell, one of which is stabilizing DNA and promoting transcription factor engagement. Another potential area of involvement for polyamines within the cholesterol pathway is the stabilization of SREBP2 binding of SRE. It has previously been shown that polyamines maintain the estrogen receptor elements (ERE) in the correct motif to allow for estrogen receptor (ER) to bind, and a lack of polyamines decreased the ability of ER to bind to EREs. Polyamines also play a role in protease function. Our lab previously demonstrated that both of CVB3’s proteases develop mutations in response to polyamine depletion, making them resistant to the lack of polyamines, suggesting a role for polyamines in protease activity. SREBP2 processing requires two proteases, S1P and S2P, for its maturation. It is unclear whether SREBP2 cleavage or S1P/S2P protease activity relies on polyamines. It is unlikely that hypusine-eIF5A is affecting another protein upstream of SREBP2 as its expression levels do not significantly change with DFMO treatment.

eIF5A is the only enzyme within the eukaryotic cell to be hypusinated, and it supports the translation of diverse cellular proteins. Prior work showed that Ebolavirus relies on hypusination specifically for the translation of VP35, a viral transactivator that facilitates viral gene expression. Other work showed that inhibitors of hypusination reduce translation of retroviruses, including HIV-1. The direct roles of eIF5A hypusination in viral protein synthesis remain to be fully explored for many viruses, including CVB3. However, hypusinated eIF5A’s roles in cellular translation also affect viral replication, as seen here. Thus, hypusinated eIF5A is an important drug target due to its potential direct and indirect effects on virus replication. Additional exploration of mechanisms connecting hypusinated eIF5A to viral and cellular factors involved in infection will further illuminate how this molecule and polyamines support the replication of diverse viruses.

## Acknowledgments

We thank The University of Chicago Genomics Facility (RRID:SCR_019196) especially Dr. Pieter Faber, for assistance with transcriptomic analyses. This work was supported by R35GM138199 from NIGMS (BCM), T32 AI007508 (MRF), and the W.M. Keck Foundation (PSS). We thank Dr. Susan Uprichard for Huh7 cells.

## Materials and methods

### Cell Culture and Virus Enumeration

Cells were maintained in Dulbecco’s modified Eagle’s medium (DMEM; Life Technologies) with bovine serum (FBS; Thermo-Fischer) and penicillin-streptomycin at 37ºC and 5% CO_2_. Huh7 cells were supplemented with 10% fetal bovine serum (FBS; Thermo-Fischer). Vero cells were obtained through BEI Resources. Vero cells were supplemented with 10% new-born calf serum (NCBS; Thermo-Fischer). CVB3 (Nancy strain) was derived from the first passage of virus in Vero cells after rescue from an infectious clone. Viral stocks were maintained at -80ºC. Viral titers were enumerated as previously described^13^.

### Drug Treatments

Difluoromethylornithine (DFMO; TargetMol) was diluted to a 100mM solution in sterile water. For DFMO treatments, cells were trypsinized (Zymo Research) and reseeded with fresh medium supplemented with 2% FBS (Huh7) or 2% NCBS (Vero). Cells were treated with 1mM DFMO unless otherwise indicated. Cells were incubated with DFMO for 96h to allow for depletion of polyamines. GC7 was diluted to a 100mM solution in sterile water. Freshly seeded cells were treated with GC7 along with 500μM aminoguanidine for 16-24h. Cholesterol was diluted to 10 ^mg^/_mL_ in ethanol then added to adherent cells for 16-24h.

### RNA Purification and cDNA Synthesis

Media were cleared from cells, and Trizol reagent (Zymo Research) was added directly. Lysate was then collected, and RNA was purified through a Zymo RNA extraction kit. Purified RNA was subsequently used for cDNA synthesis using High Capacity cDNA Reverse Transcription Kits (Thermo-Fischer), according to the manufacturer’s protocol, with 10–100 ng of RNA and random hexamer primers.

### RNA Sequencing

RNA was purified and prepared as described from Huh7 cells treated for 96h with DFMO or infected for 24h with CVB3. Libraries were prepared by the University of Chicago Genomics Facility and analyzed by Illumina NovaSeq 6000. Read quality was evaluated using FastQC (v0.11.5). Adapters were trimmed in parallel to a quality trimming (bbduk, sourceforge.net/projects/bbmap/). All remaining sequences were mapped against the human reference genome build 38 with STAR (v2.5.2b)^30^. HTseq (v0.6.1) was used to count all reads for each gene and set up a read count table^31^. Differential gene expression analyses were performed using the DESeq2 Bioconductor package (v1.30.1)^32^. The default “ashr” shrinkage (v2.2-47)^33^ set up was used for our analysis. Gene set enrichment analysis (GSEA) was performed with the fgsea Bioconductor package^34^, using Hallmark gene sets downloaded from the Molecular Signatures Database^35^.

### Plaque Formation Attachment Assay

Vero cells were seeded in 6-well plates and grown to 100% confluence in DMEM with 2% NCBS and treated for 96h with the indicated concentrations of DFMO. After 96 h of DFMO treatment, cells were placed on ice and the media aspirated from the cells. 500 uL of serum free media containing 1000 PFU CVB3 was added to cells on ice for 5 min. Cells were washed 3x with PBS and then overlaid with 0.8% agarose containing DMEM with 2% NCBS. The plates were incubated at 37ºC for 2 days for plaques to develop. The cells were fixed with 4% formalin, and the plaques were visualized with crystal violet staining. For the cholesterol rescue, cells were washed 3x with PBS before infecting with CVB3.

### qPCR Gene Expression Assay

Huh7 cells were seeded at 4 × 10^4^ cells per well in 24-well plates in DMEM with 2% FBS. Cells were treated with varying concentrations of DFMO for 96 h. After 96 h, the media was aspirated off cells, washed 1x with PBS, and then, 200 uL of Trizol was added to the cells. The RNA was extracted with the Zymo RNA extraction kit, converted to cDNA, and quantified by real-time PCR with SYBR Green (DotScientific) using the one-step protocol QuantStudio 3 (ThermoFisher Scientific). Relative expression was calculated using the ΔΔC^T^ method, normalized to the β-actin qRT-PCR control, and calculated as the fraction of the untreated samples. Primers were verified for linearity using 8-fold serial diluted cDNA and checked for specificity via melt curve analysis. The primer sequences are as follows: HMGCR, (F) 5’-GAG ACA GGG ATA AAC CGA GAA AG-3’ and (R): 5’-GGA GGA GTT ACC AAC CAC AAA-3’; HMGCS, (F): 5’-CCT GCC AAG AAA GTA CCA AGA-3’ and (R): 5’-GTC TTG CAC CTC ACA GAG TAT C-3’; MVD (F): 5’-TGG TTC TGC CCA TCA ACT C-3’ and (R): 5’-GGT GAA GTC CTT GCT GAT GA-3’; SREBP2 (F): 5’-CTG TAG CGT CTT GAT TCT CTC C-3’ and (R): 5’-CCT GGC TGT CCT GTG TAA TAA-3’.

### Western Blot

Samples were collected with Bolt LDS Buffer and Bolt Reducing Agent (Invitrogen, Waltham, MA, USA) and run on polyacrylamide gels. Gels were transferred using the iBlot 2 Gel Transfer Device (Invitrogen). Membranes were blocked with 5% BSA in TBST then probed with primary antibodies for HMGCR (Ms mAb, 1:000, abcam), MVD (1:1000, Santa Cruz), SREBP2 (Gt pAb, 1:1000, R&D Systems), Hypusine (Rb pAb, 1:2000, EMD Millipore) orβ-actin (Ms mAb, 1:1000, proteintech) overnight at 4ºC. Membranes were then washed 3x in TBST followed by 1h incubation of in secondary antibody (GtαMs/GtαRb/DonkeyαGt HRP, 1:15000, Jackson Labs). After 3 additional washes in TBST, membranes were treated with SuperSignal West Pico PLUS Chemiluminescent Substrate (ThermoFisher Scientific) and visualized on Fluorchem E imager (Protein Simple, San Jose, CA, USA). Quantification of western blots were done by using ImageJ and normalizing to NT and relative to actin density.

### Intracellular Cholesterol Abundance Assay

Huh7 cells were plated at a density of 5000 cells/well in a 96 well plate in DMEM with 2% FBS. Cells were treated with DFMO for 96h or after 72h, treated with DEF or GC7 for 24h. The following day, the media was removed from cells followed by a PBS wash. To measure total intracellular cholesterol abundance, we used the Cholesterol/Cholesterol Ester-Glo Assay™ (Promega) in accordance to manufacturer’s protocol.

### SRE promoter luciferase

Complimentary primers were made containing SRE consensus sequence were ordered flanked by SfiI cut site overhangs (FWD: 5’-CGGCC ATCACCCCAC GGCCTCGG-3’; REV 3’-GCCGCCGG TAGTGGGGTG CCGGA-5’). Primers were phosphorylated and annealed at 37ºC for 30 minutes then 95ºC for 5 minutes and were allowed to cool to 25ºC. pGL4.10 (Promega) was digested with SfiI in Fast Digest buffer for 15 min at 50ºC. The cut plasmid was ran through DNA clean up kit. The annealed primers were then ligated into the cut plasmid using T4 ligase followed by transformation into chemical competent E. Coli. Colonies were picked and grown up followed by sequencing to confirm the SRE sequence was present.

### Promoter Luciferase Assay

Huh7 cells were plated in a 96 well plate with 2% FBS DMEM then treated with 1 mM DFMO for 96 h or after 96 h, treated with 500 uM GC7. Cells were transfected with SRE-pGL4.10, 5’ HMGCS-Fluc (addgene #60444), or pLDLR-Luc (addgene #14940) after cells had been plated for 96 h. All cells were transfected with the renilla control plasmid siCheck (Promega). 100 ng of plasmid were transfected with LipoD293 according to manufacture’s protocol. 24 h after transfection, media was removed followed by one wash with PBS. Cells were then lysed with gentle lysis buffer for 15 min.

### Statistical analysis

Prism 6 (GraphPad) was used to generate graphs and perform statistical analysis. For all analyses, a two-tailed Student’s *t* test was used to compare groups, unless otherwise noted.

## References

1. Mounce, B. C., Olsen, M. E., Vignuzzi, M. & Connor, J. H. Polyamines and Their Role in Virus Infection. Microbiol. Mol. Biol. Rev. MMBR 81, (2017).

2. Burri, C. & Brun, R. Eflornithine for the treatment of human African trypanosomiasis. Parasitol. Res. 90 Supp 1, S49–52 (2003).

3. Park, M. H., Cooper, H. L. & Folk, J. E. Identification of hypusine, an unusual amino acid, in a protein from human lymphocytes and of spermidine as its biosynthetic precursor. Proc. Natl. Acad. Sci. U. S. A. 78, 2869–2873 (1981).

4. Schuller, A. P., Wu, C. C.-C., Dever, T. E., Buskirk, A. R. & Green, R. eIF5A Functions Globally in Translation Elongation and Termination. Mol. Cell 66, 194-205.e5 (2017).

5. Gutierrez, E. et al. eIF5A Promotes Translation of Polyproline Motifs. Mol. Cell 51, 35–45 (2013).

6. D’Agostino, M. et al. Insights Into the Binding Mechanism of GC7 to Deoxyhypusine Synthase in Sulfolobus solfataricus: A Thermophilic Model for the Design of New Hypusination Inhibitors. Front. Chem. 8, 1170 (2020).

7. Park, M. H., Joe, Y. A., Kang, K. R., Lee, Y. B. & Wolff, E. C. The polyamine-derived amino acid hypusine: its post-translational formation in eIF-5A and its role in cell proliferation. Amino Acids 10, 109–121 (1996).

8. Mounce, B. C. et al. Chikungunya Virus Overcomes Polyamine Depletion by Mutation of nsP1 and the Opal Stop Codon To Confer Enhanced Replication and Fitness. J. Virol. 91, (2017).

9. Olsen, M. E., Cressey, T. N., Mühlberger, E. & Connor, J. H. Differential Mechanisms for the Involvement of Polyamines and Hypusinated eIF5A in Ebola Virus Gene Expression. J. Virol. 92, (2018).

10. Mastrodomenico, V. et al. Polyamine depletion inhibits bunyavirus infection via generation of noninfectious interfering virions. J. Virol. JVI.00530-19 (2019) doi:10.1128/JVI.00530-19.

11. Firpo, M. R. et al. Targeting Polyamines Inhibits Coronavirus Infection by Reducing Cellular Attachment and Entry. ACS Infect. Dis. acsinfecdis.0c00491 (2020) doi:10.1021/acsinfecdis.0c00491.

12. Dial, C. N., Tate, P. M., Kicmal, T. M. & Mounce, B. C. Coxsackievirus B3 Responds to Polyamine Depletion via Enhancement of 2A and 3C Protease Activity. Viruses 11, 403 (2019).

13. Kicmal, T. M., Tate, P. M., Dial, C. N., Esin, J. J. & Mounce, B. C. Polyamine depletion abrogates enterovirus cellular attachment. J. Virol. JVI.01054-19 (2019) doi:10.1128/JVI.01054-19.

14. Hulsebosch, B. M. & Mounce, B. C. Polyamine Analog Diethylnorspermidine Restricts Coxsackievirus B3 and Is Overcome by 2A Protease Mutation In Vitro. Viruses 13, 310 (2021).

15. Pons-Salort, M., Parker, E. P. K. & Grassly, N. C. The epidemiology of non-polio enteroviruses: recent advances and outstanding questions. Curr. Opin. Infect. Dis. 28, 479–487 (2015).

16. Massilamany, C., Gangaplara, A. & Reddy, J. Intricacies of cardiac damage in coxsackievirus B3 infection: Implications for therapy. Int. J. Cardiol. 177, 330–339 (2014).

17. Chapman, N. M. & Kim, K.-S. Persistent Coxsackievirus Infection: Enterovirus Persistence in Chronic Myocarditis and Dilated Cardiomyopathy. in Group B Coxsackieviruses 275–292 (Springer, Berlin, Heidelberg, 2008). doi:10.1007/978-3-540-75546-3_13.

18. Coxsackievirus B3 replication and pathogenesis | Future Microbiology. https://www.futuremedicine.com/doi/full/10.2217/fmb.15.5.

19. Archard, L. C. et al. Molecular probes for detection of persisting enterovirus infection of human heart and their prognostic value. Eur. Heart J. 12, 56–59 (1991).

20. Mounce, B. C. et al. Inhibition of Polyamine Biosynthesis Is a Broad-Spectrum Strategy against RNA Viruses. J. Virol. 90, 9683–9692 (2016).

21. Danthi, P. & Chow, M. Cholesterol Removal by Methyl-β-Cyclodextrin Inhibits Poliovirus Entry. J. Virol. 78, 33–41 (2004).

22. Stuart, A. D., Eustace, H. E., McKee, T. A. & Brown, T. D. K. A Novel Cell Entry Pathway for a DAF-Using Human Enterovirus Is Dependent on Lipid Rafts. J. Virol. 76, 9307–9322 (2002).

23. Ilnytska, O. et al. Enteroviruses harness the cellular endocytic machinery to remodel the host cell cholesterol landscape for effective viral replication. Cell Host Microbe 14, 281–293 (2013).

24. 6.35 Cholesterol Synthesis | Nutrition Flexbook. https://courses.lumenlearning.com/suny-nutrition/chapter/6-35-cholesterol-synthesis/.

25. Hua, X. et al. SREBP-2, a second basic-helix-loop-helix-leucine zipper protein that stimulates transcription by binding to a sterol regulatory element. Proc. Natl. Acad. Sci. 90, 11603–11607 (1993).

26. Luo, J., Yang, H. & Song, B.-L. Mechanisms and regulation of cholesterol homeostasis. Nat. Rev. Mol. Cell Biol. 21, 225–245 (2020).

27. Pirinen, E. et al. Activated polyamine catabolism leads to low cholesterol levels by enhancing bile acid synthesis. Amino Acids 38, 549–560 (2010).

28. Brown, A. P., Morrissey, R. L., Crowell, J. A. & Levine, B. S. Difluoromethylornithine in combination with tamoxifen in female rats: 13-week oral toxicity study. Cancer Chemother. Pharmacol. 44, 475–483 (1999).

29. Rossi, D. et al. eIF5A has a function in the cotranslational translocation of proteins into the ER. Amino Acids 46, 645–653 (2014).

30. Dobin, A. et al. STAR: ultrafast universal RNA-seq aligner. Bioinformatics 29, 15–21 (2013).

31. Anders, S., Pyl, P. T. & Huber, W. HTSeq—a Python framework to work with high- throughput sequencing data. Bioinformatics 31, 166–169 (2015).

32. Love, M. I., Huber, W. & Anders, S. Moderated estimation of fold change and dispersion for RNA-seq data with DESeq2. Genome Biol. 15, 550 (2014).

33. Stephens, M. False discovery rates: a new deal. Biostatistics 18, 275–294 (2017).

34. Korotkevich, G. et al. Fast gene set enrichment analysis. 060012 https://www.biorxiv.org/content/10.1101/060012v3 (2021) doi:10.1101/060012.

35. Liberzon, A. et al. Molecular signatures database (MSigDB) 3.0. Bioinformatics 27, 1739–1740 (2011).

